# Definition of naturally processed peptides reveals convergent presentation of autoantigenic topoisomerase-I epitopes in scleroderma

**DOI:** 10.1101/798249

**Authors:** Eleni Tiniakou, Andrea Fava, Zsuzsanna H. McMahan, Tara Guhr, Robert N. O’Meally, Ami A. Shah, Fredrick M. Wigley, Robert N. Cole, Francesco Boin, Erika Darrah

## Abstract

Disease-associated *HLA-DRB1* alleles are thought to confer risk of developing autoimmunity by favoring the presentation of select autoantigenic epitopes. However, identification of these epitopes and the principles governing their presentation has been hindered by the imprecision of currently available methods, which cannot fully recapitulate the complexity of human pathophysiology. We present a natural antigen processing assay (NAPA), which overcomes these limitations by studying the presentation of autoantigenic CD4+ T cell epitopes by monocyte-derived dendritic cells (mo-DCs) from patients. We applied this strategy to study the processing and presentation of topoisomerase-1 (TOP1), a prevalent autoantigen in scleroderma that is associated with lung fibrosis and high mortality. We found that a common set of 10 epitopes was presented by mo-DCs from patients with diverse HLA-DR variants, including those not previously associated with the disease. Sequence analysis revealed a shared peptide-binding motif within the HLA-DR peptide binding grooves of patients who developed anti-TOP1 autoantibodies. In addition, a subset of naturally presented TOP1 peptides were characterized by immunological promiscuity, as they could bind to diverse HLA-DR peptide binding grooves. NAPA epitopes were immunorelevant: they could stimulate autoreactive CD4+ T cells in patients, and the number of epitopes recognized correlated with lung disease severity. These findings mechanistically implicate presentation of a convergent set of TOP1 epitopes in the development of scleroderma lung disease. Precise identification of autoantigenic epitopes is key to understanding the primordial mechanisms for the loss of tolerance, studying disease-propagating autoreactive T cells, and developing antigen-specific immunotherapy.

**One Sentence Summary:** Use of a novel natural antigen processing assay reveals a mechanism for the presentation of shared CD4+ T cell epitopes of topoisomerase-I in immunogenetically diverse patients with scleroderma.

## Introduction

Targeting of protein autoantigens by the adaptive immune system is a hallmark feature of autoimmune diseases. As fundamental regulators of cellular and humoral immunity, CD4+ T cells specific for self-proteins are implicated as central pathogenic drivers of these diseases(1). This is highlighted by high-titer autoantibodies of the IgG subclass, indicative of CD4+ T helper cell involvement, and strong genetic association with specific *HLA-DRβ1* alleles observed in many autoimmune disorders. Despite the recognized importance of autoantigen-specific CD4+ T cells in autoimmune disease pathogenesis, our current knowledge of autoreactive T cell epitope specificity is limited, and even less is known about the mechanisms that govern the presentation of autoantigenic epitopes by disease-associated HLA-DR molecules. This lack of understanding is largely due to: 1) the complexity of studying disease in highly heterogeneous human populations; 2) limited access to patient biospecimens; and 3) the low sensitivity and high cost of currently available methods for studying autoreactive T cells in humans. This has led to the reliance on instructive animal models and the widespread use of *in silico* prediction algorithms or overlapping peptide libraries for defining candidate autoreactive CD4+ T cell epitopes(2, 3). While informative, these methods cannot recapitulate the nuances of human pathophysiology, highlighting the need to approach autoantigenic CD4+ T cell epitope discovery by studying antigen processing directly in patients.

Scleroderma, also called systemic sclerosis, is a complex autoimmune rheumatic disease of unknown etiology, characterized by widespread vasculopathy, fibrosis of the skin and internal organs(4), and immunological derangements, including the production of autoantibodies to specific nuclear antigens(5). No existing animal model fully recapitulates the intricate immunopathology of this condition. Anti-topoisomerase-I (TOP1) autoantibodies (ATA; also known as anti-Scl-70) are present in 20-45% of patients with scleroderma and are associated with diffuse skin involvement, pulmonary fibrosis and high mortality(6). ATA production in patients with scleroderma is associated with specific *HLA-DRB1* alleles, implicating CD4+ T helper cells in driving immune responses to TOP1. TOP1-specific CD4+ T cells have been detected in the peripheral blood of patients with scleroderma(7–10), are HLA-DR restricted, and their frequency is associated with the presence, severity, and progression of lung fibrosis(11). Previous attempts to define TOP1 epitopes in patients with scleroderma using overlapping peptide/protein fragment libraries or peptides derived from *in silico* prediction have resulted in the identification of scattered epitopes with unspecified biological and clinical significance(7–10). Therefore, although ATA-associated *HLA-DRB1* alleles have been defined and TOP-I-specific CD4+ T cells have been detected in ATA-positive patients, the identification of the TOP1 epitopes driving the autoimmune process has remained elusive.

To overcome current limitations related to the identification and study of autoreactive T cell epitopes, we developed a method for identifying the repertoire of naturally processed TOP1 peptides presented by antigen presenting cells (APCs) obtained from patients with scleroderma. This approach has innovative features compared to traditional epitope-discovery methods, as it harnesses the cellular MHC class II antigen processing machinery to identify naturally presented peptides, without requiring pre-defined knowledge about the target peptide length or HLA status. Our analysis of naturally presented TOP1 peptides has revealed unexpected insights into the pathogenesis of scleroderma and identified a discrete set of clinically relevant TOP1-specific CD4+ T cell epitopes that may inform future development of antigen-specific diagnostic and therapeutic tools.

## Results

### Identification of naturally processed TOP1 peptides

We developed a natural antigen processing assay (NAPA) to identify TOP1 peptides presented by MHC class II molecules expressed on antigen presenting cells (APC) from patients with scleroderma. As a source of patient-specific APCs, monocytes were isolated from the peripheral blood of 6 ATA-positive patients with scleroderma and cultured in the presence of GM-CSF and IL-4 to generate monocyte-derived dendritic cells (mo-DCs). Mo-DCs were then pulsed with the whole TOP1 protein to allow for internalization and processing. HLA-DR-peptide complexes were isolated by immunoprecipitation (experimental workflow outlined in Fig. 1). The HLA-DR bound peptides were then eluted, separated by size exclusion, and sequenced by mass spectrometry.

**Fig. 1.**
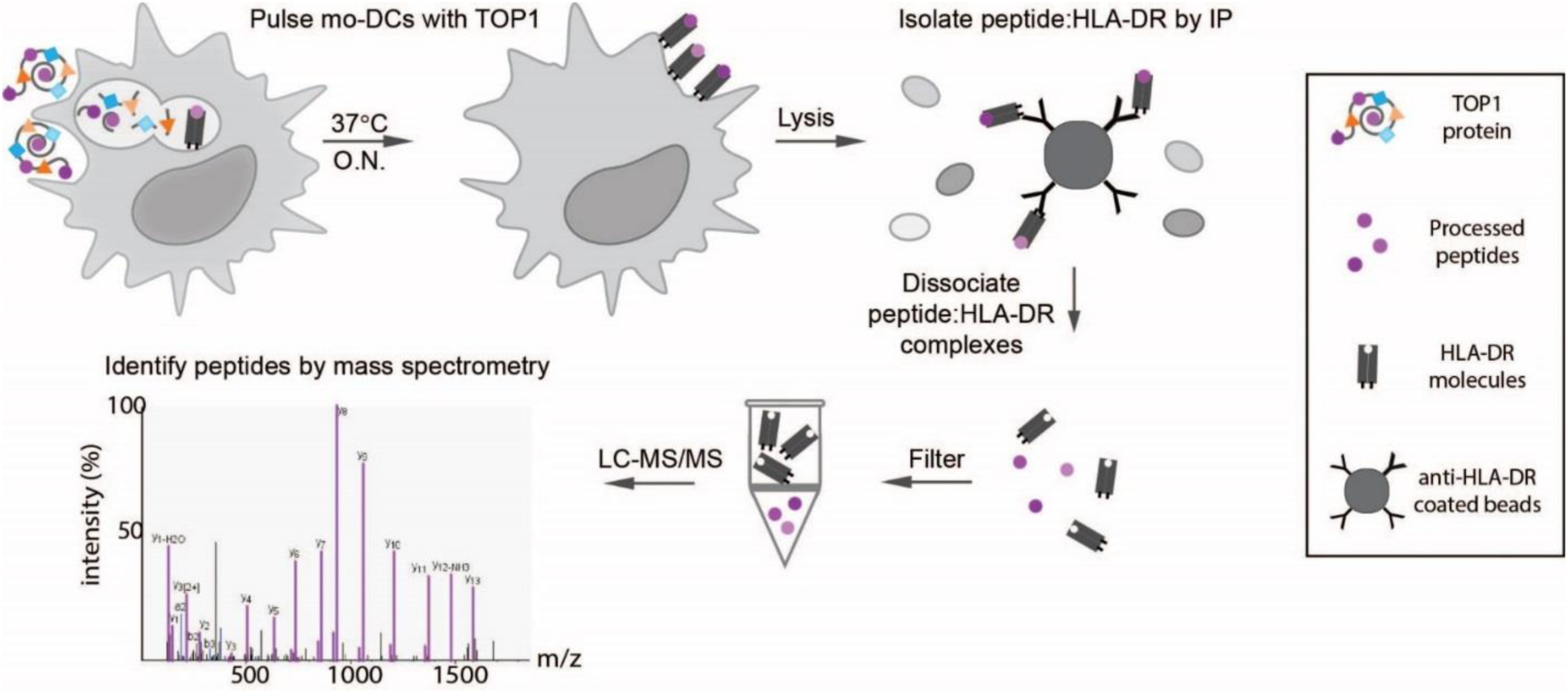
Natural Antigen Processing Assay (NAPA). MoDCs from 6 randomly selected donors with ATA-positive scleroderma were pulsed with TOP1 protein over-night (O.N.) at 37°C. Cells were then lysed and the peptide:HLA-DR complexes isolated using magnetic beads coated with anti-HLA-DR antibodies (clone L243). The peptides were dissociated from the HLA-DR molecules using 0.1% TFA and separated from HLA-DR using a 10kDa spin filter prior to sequencing by liquid chromatography with tandem mass spectrometry (LC-MS/MS).

NAPA identified hundreds of naturally processed peptides, deriving from TOP1 as well as from other self-proteins (Fig. 2 and Table S1). The median length of these peptides was 15 amino acids. Clustering into nested sets of variable length peptides around a common core sequence was common (Table S1). These are features typical of naturally processed MHC class II peptides, which tend to be 13-25 amino acids long and exist in nested sets (12, 13). As expected, pathway analysis of the naturally processed peptides showed that the majority of these peptides were derived from proteins involved in antigen processing (Fig. 2B). Further analysis of the peptides presented between individuals revealed an enrichment of peptides from proteins expected to traffic through the secretory pathway, where MHC class II antigen processing occurs (Fig. 2C). These including membrane-associated (e.g. ITGAM and MRC1) and secreted proteins (e.g. TGFBI and TIMP3), as well as other MHC molecules. Importantly, the only antigen commonly presented by all 6 patients was the pulsed antigen TOP1, demonstrating the ability of NAPA to detect peptides from exogenous proteins. The specificity of NAPA for detecting HLA-DR-bound peptides, including those derived from TOP1, was confirmed using unpulsed mo-DCs and isotype control antibodies for immunoprecipitation (Tables S2 and S3). In these experimental conditions, no TOP1 peptides were detected.

**Fig. 2.**
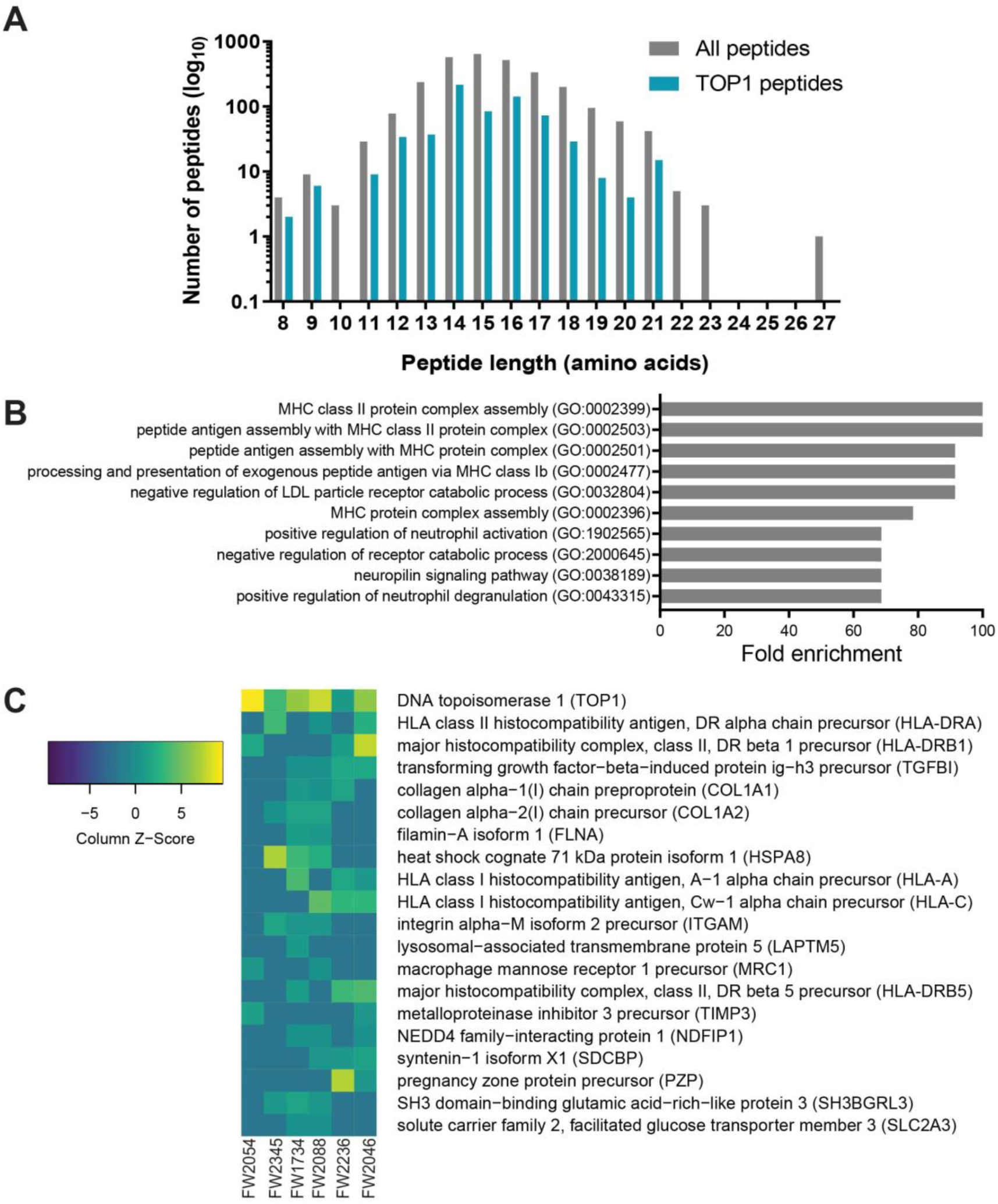
Characteristics of the naturally processed peptides isolated from NAPA. **(A)** Histogram demonstrating the distribution of naturally presented peptides based on the amino acid length of all peptides(grey) and TOP1 peptides (teal) isolated from 6 patients with ATA-positive SSc. **(B)** The “GO biological processes complete” annotation set was queried to identify pathways that were overrepresented in the naturally processed peptides relative to the expected enrichment in the human genome. FDR and Fisher’s exact p-value for each pathway was calculated. The 10 most highly enriched biological processes are shown with FDR <0.03 and Fisher’s exact p-value <0.001. (C) Heatmap illustrating the frequency of the most commonly naturally presented peptides between all patients.

Remarkably, we observed a substantial overlap in TOP1 peptides presented by ATA-positive scleroderma patients, with all subjects presenting peptides originating from only 10 discrete 13-23 amino acid long regions of the TOP1 molecule (Fig. 3A). The median number of putative regions presented by each individual patient was 4 (range 1-7), and 70% were presented by at least two subjects. The 10 regions from which TOP1 peptides were presented are from diverse domains in the TOP1 protein and largely reside within structural elements, most dominantly alpha helices flanked by unstructured loops on the surface of the TOP1 protein (Fig.3A and B). The localization of putative TOP1 epitopes within structural elements adjacent to flexible loops is similar to what has been observed for CD4+ T cell epitopes derived from other infectious and self-antigens(14, 15). Using NAPA, we successfully identified a common set of TOP1 peptides derived from the whole TOP1 protein, which were naturally processed and presented by HLA-DR molecules in mo-DCs generated *ex vivo* from patients with ATA-positive scleroderma.

**Fig. 3.**
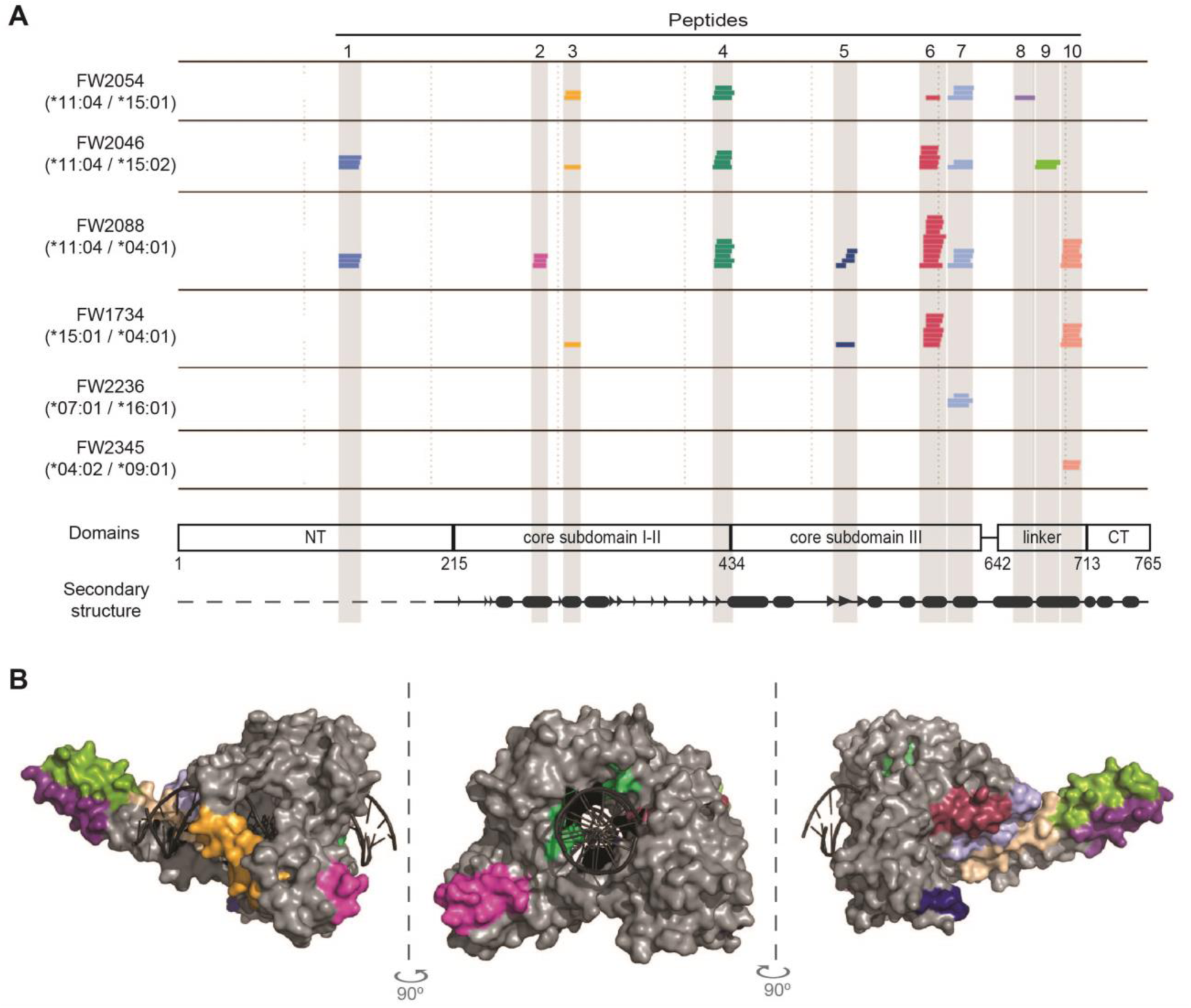
A common set of naturally processed TOP1 peptides was presented by scleroderma patients with diverse HLA-DR variants. **(A)** The *HLA-DRB1* haplotype for each patient is given and each colored line represents a unique TOP1 peptide sequence identified by mass spectrometry. The location of the TOP1 peptides within the linear TOP1 protein sequence is indicated. TOP1 protein domains and secondary structure elements are shown (β-strands and α-helices are represented by arrow heads and ovals, respectively). **(B)** The location of the naturally processed TOP1 peptides within the three-dimensional crystal structure of the TOP1 molecule bound to DNA (1K4T) is indicated. The peptides are colored-coded and correspond to the colors used in panel A.

### Shared motifs present in HLA-DRβ chains of patients with ATA-positive scleroderma

We next performed HLA immunogenetic analysis to determine if the presentation of common TOP1 peptides identified in our patients was due to the presence of shared *HLA-DRB1* alleles. Despite the known association of ATA with specific *HLA-DRB1* alleles (i.e. *HLA-DRB1*\*11:01, *11:04, *08:02, *08:04, or *15:02)(16), only 3 subjects (50%) harbored a previously defined risk allele. While these 3 patients concurrently presented peptides from more regions of the TOP1 molecule compared to those who did not exhibit a known ATA-associated *HLA-DRB1* allele (median 6 vs. 1), common peptides were presented regardless of the allele status.

Given this unexpected ability of diverse HLA-DR variants to present common TOP1 peptides, we sought to determine if similar features were present in the HLA-DRβ chains of ATA-positive scleroderma patients. To explore this, we performed amino acid sequence alignment of the HLA-DRβ variants linked to the development of ATA-positive scleroderma(16) and compared them to the HLA-DRβ chains in these patients. This analysis revealed two conserved motifs within the DRβ-chain encoded for by the *HLA-DRB1* alleles most strongly linked to ATA(17–20) (*HLA-DRB1*\*08:02, *08:04, *11:01, *11:04): ^9^E-Y-S-T-S/G-E^14^ and ^67^F-L-E-D-R-R-A-A/L^74^ (Fig. 4A). HLA-DRβ1*15:02, associated with ATA in Asian populations(16), had a similar ^67^I-L-E-Q-A-R-A-A^74^ motif but a distinct sequence at position 9-14 (^9^W-Q-P-K-R-E^14^) (Fig. 4A). Importantly, these two motifs contain key peptide contact residues at positions 9, 11, 13, 14 and 67, 70, 71, 74 respectively (Fig. 4B).

**Fig. 4.**
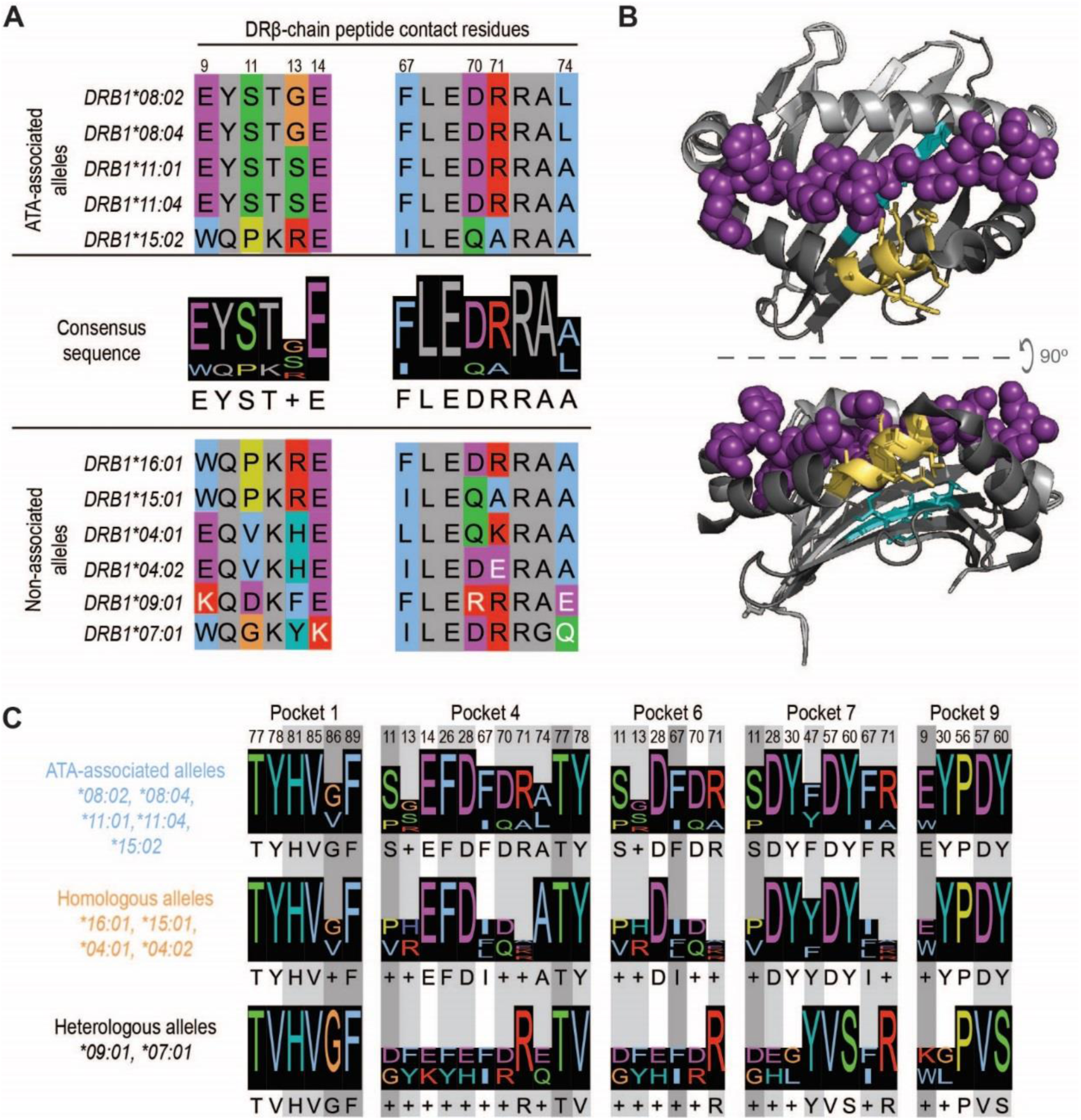
Presentation of common TOP1 peptides linked to shared ATA-associated motifs in the HLA-DRβ chain. **(A)** Amino acid sequence alignment of ATA-associated (top panel) and non-associated alleles (bottom panel) is shown. The two consensus sequences identified in the peptide-binding groove of ATA-associated alleles are shown (middle panel). Peptide contact residues in the DRβ chain are shown (hydrophobic=blue, negative=purple, positive=red, polar=green, glycine=orange, prolines=yellow, and aromatic=cyan) and their position within the sequence is numbered, while intervening residues are shown in gray. The white text indicates non-conserved residues. (B) A depiction of HLA-DR11 bound to peptide (6CPN) is shown with the peptide binding groove: DR α-chain residues (light gray), DR β-chain (dark gray) and peptide (magenta). The ^9^E-Y-S-T-S/G^13^ and ^67^F-L-E-D-R-R-A-A/L^74^ motifs are shown in cyan and yellow, respectively. **(C)** Peptide contact residues for each MHC binding pocket and the relevant consensus sequence for ATA-associated (light blue), homologous (orange) and heterologous (black) alleles are represented. Different shades of gray indicate the importance of each peptide residue to peptide binding, with dark gray being the most critical.

Analysis of the HLA-DRβ chains present in patients who did not have previously defined ATA-associated *HLA-DRB1* alleles, but were able to present common TOP1 peptides, revealed several conserved features. HLA-DRβ1*15:01 and *16:01 contained motifs at positions 9-14 and 67-74 that were identical to known ATA-associated alleles (Fig. 4A). HLA-DRβ1*04:01 and *04:02 were less similar, but had conserved amino acids at known peptide contact positions (i.e. 9, 14, and 67, 70, 74). A more detailed analysis of β-chain residues that comprise the five major peptide binding pockets in the HLA-DR groove (denoted as P1, P4, P6, P7, and P9) confirmed that ATA-associated and homologous alleles have highly similar peptide binding pockets (Fig. 4C). Interestingly, HLA-DRβ1*09:01 and *07:01 were divergent from ATA-associated alleles at several critical peptide contact residues (Fig. 4, A and C). HLA-DRβ1*07:01 has been reported to be protective against scleroderma development(21). While all the ATA-positive patients in our study possessed at least one allele that contained ATA-associated motifs, those who harbored HLA-DRβ1*09:01 and *07:01 presented the fewest number of TOP1 peptides (Fig. 3A). These data reveal shared features in the peptide binding grooves of seemingly unrelated HLA-DRβ1 variants present in ATA-positive patients affected by scleroderma, highlighting their ability to present a common set of TOP1 peptides.

### Promiscuous binding of TOP1 peptides to multiple HLA-DR variants

To examine the intermolecular interactions between the commonly presented TOP1 peptides identified using NAPA and the HLA-DR variants detected in these patients, we used the NetMHCII 2.3 algorithm to predict the peptide binding affinity and register(22). Since all of the patients were heterozygous at the *HLA-DRB1* locus, we examined the predicted binding of peptides to both HLA-DR variants present in a given individual. This analysis revealed peptides that could be categorized into two distinct groups, termed group A and B, based on their predicted binding register (Fig. 5). Group A peptides (peptides 1, 2, 8, 9, and 10) were predicted to bind in the same register to all associated HLA-DR variants. This is in contrast to group B peptides, which were predicted to bind in different registers to multiple different HLA-DR variants. Group A peptides were also predicted to bind with significantly lower affinity to their respective HLA-DR molecules than Group B peptides (peptides 3, 4, 5, 6, and 7) (4421 ± 1184 vs. 535 ± 276 nM; p=0.013), and were presented by fewer individuals than group B peptides (median 1 vs. 3). This suggests that group A peptides are relatively weak-binding peptides that contain a single core epitope (e.g. LMKLEVQAT in peptide 10), whose anchor residues interact with shared features present in the HLA-DR binding grooves of patients with ATA. In contrast, group B peptides appear to contain multiple distinct epitopes, each possessing a unique set of anchor residues that enable high-affinity interaction with a more diverse set of HLA-DR binding pockets (e.g. LSYNRANRA, ILSYNRANR, and YNRANRAVA in peptide 7). Taken together, these data support a model of convergent TOP1 peptide presentation in patients with scleroderma driven by shared peptide-binding motifs in the HLA-DR groove and a promiscuous group of TOP1 peptides.

**Figure 5.**
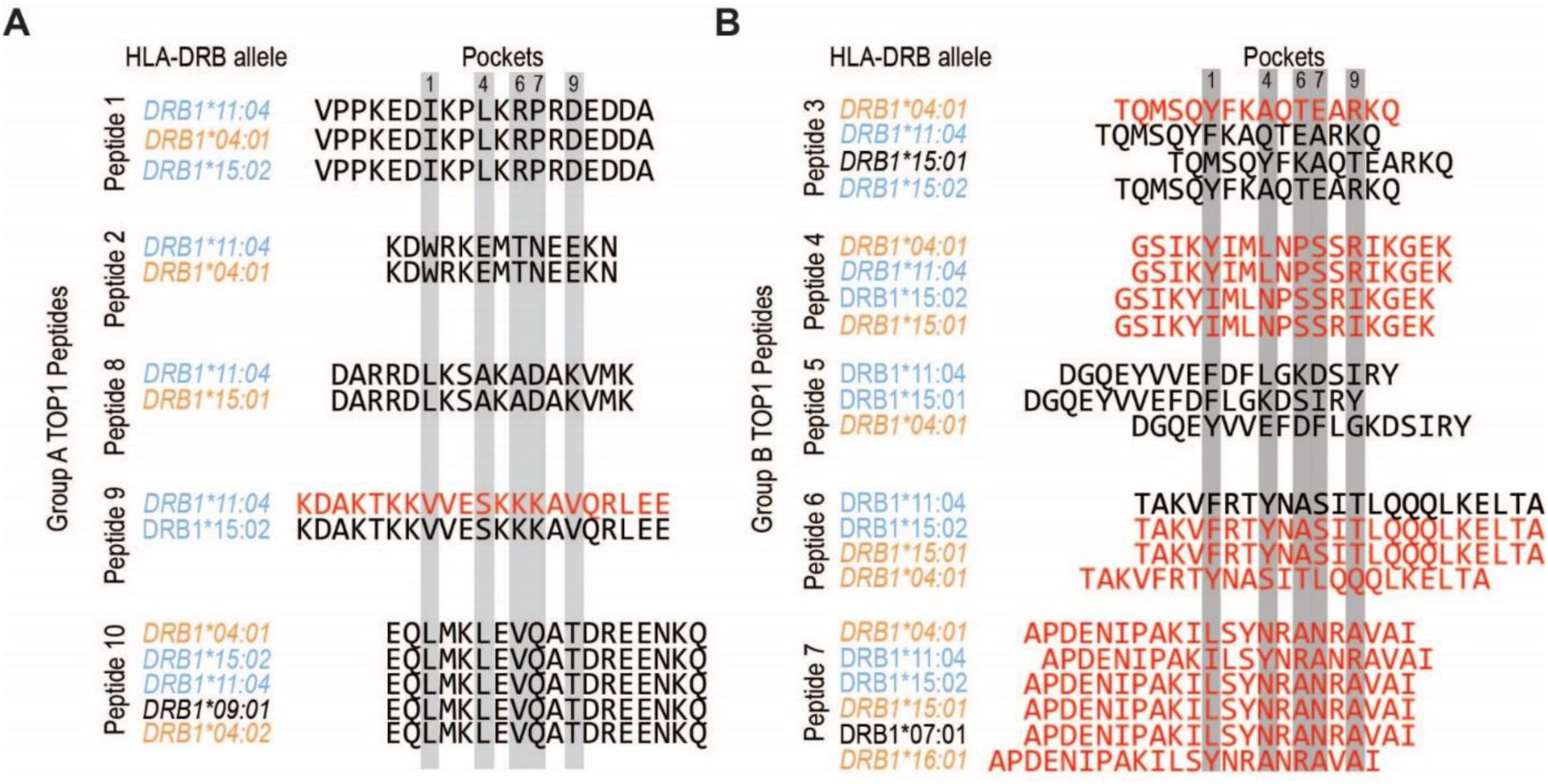
Allele-specific TOP1 peptide binding registers may contribute to promiscuous presentation by diverse HLA-DR variants. **(A)** Group A TOP1 peptides (peptides 1, 2, 8, 9 and 10) and **(B)** group B TOP1 peptides (peptides 3, 4, 5, 6 and 7) are aligned according to their predicted binding registers as per NetMHCII algorithm with relevant anchor residues highlighted in gray. The *HLA-DRB1* alleles are color-coded as ATA-associated (light blue), homologous (orange) and heterologous (black) alleles, and ordered from highest to lowest predicted affinity for each peptide. TOP1 peptides with a predicted binding affinity of <500 nM are indicated with red text, and are expected to represent moderate to high affinity peptides(22).

### Robust recognition of commonly presented TOP1 epitopes by CD4+ T cells in scleroderma

The discovery of a shared mechanism for the presentation of a core set of TOP1 peptides suggests that these epitopes may be central to the disease process in patients with scleroderma. To determine if these epitopes are immunologically relevant and recognized in ATA-positive patients, we synthesized 10 peptides corresponding to the putative TOP1 epitopes identified using NAPA (Fig. 3A; Table 1) and examined their ability to stimulate CD4+ T cells from 11 ATA-positive and 11 ATA-negative scleroderma patients (clinical variables are shown in Table S4). CD4+ T cell activation, as measured by upregulation of the early activation marker CD154(11, 23), was observed in response to at least one TOP1 peptide in 73% (8/11) of ATA-positive patients compared to 27% (3/11) of ATA-negative individuals (Figure 6A, Table S5). ATA-positive patients also responded concurrently to significantly more TOP1 peptides (median 2 vs. 0; p=0.02) (Fig. 6B), particularly ATA-positive subjects with known disease-associated *HLA-DRB1* alleles (median 7 vs. 1.5). Recognition of group A and B peptides was observed regardless of ATA-associated allele status. For 6 out of the 11 ATA-positive patients (54%), the combined frequency of CD154+ cells observed in response to the individual TOP1 peptides (Fig. S1) was comparable to that measured in response to the whole TOP1 protein, suggesting the immunodominance of these 10 naturally processed TOP1 peptides in these individuals. Robust recognition of this core set of TOP1 peptides in immunogenetically diverse ATA-positive patients, supports our observation that features on both sides of the peptide-MHC interaction contribute to the development of a meaningful anti-TOP1 immune responses in patients with scleroderma.

**Table 1.**
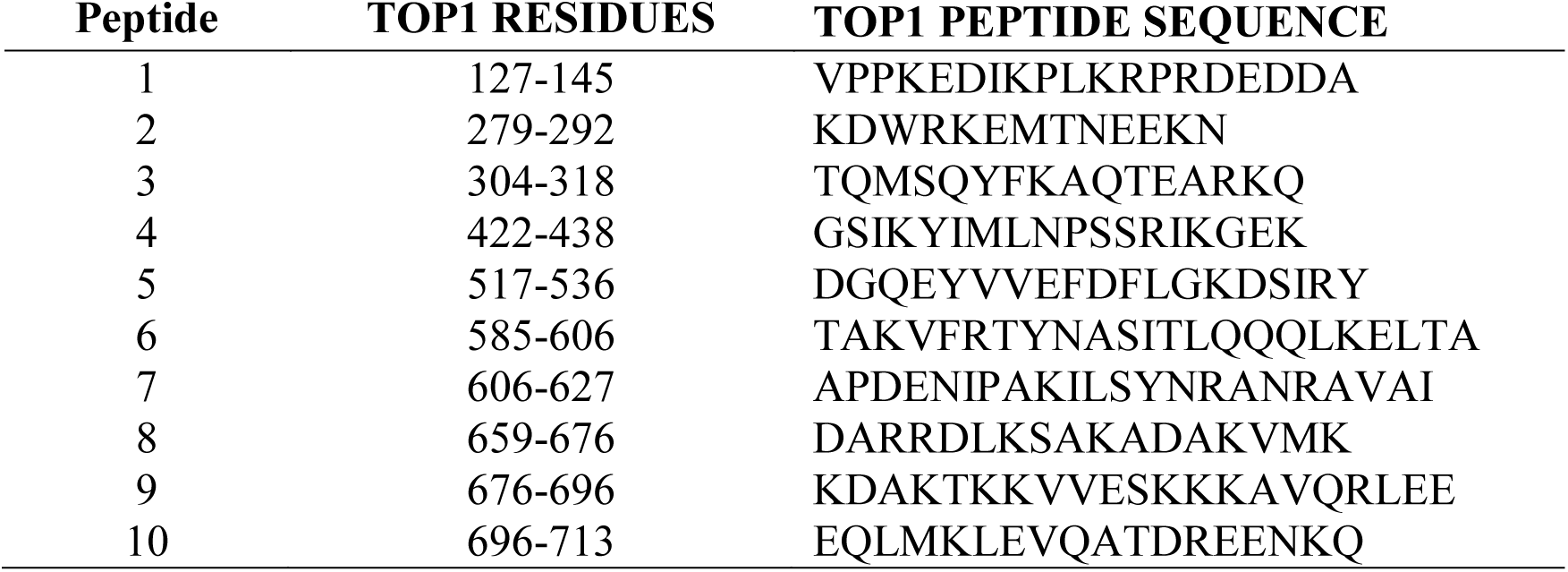
TOP1 peptides corresponding to the TOP1 regions identified by NAPA

**Fig. 6.**
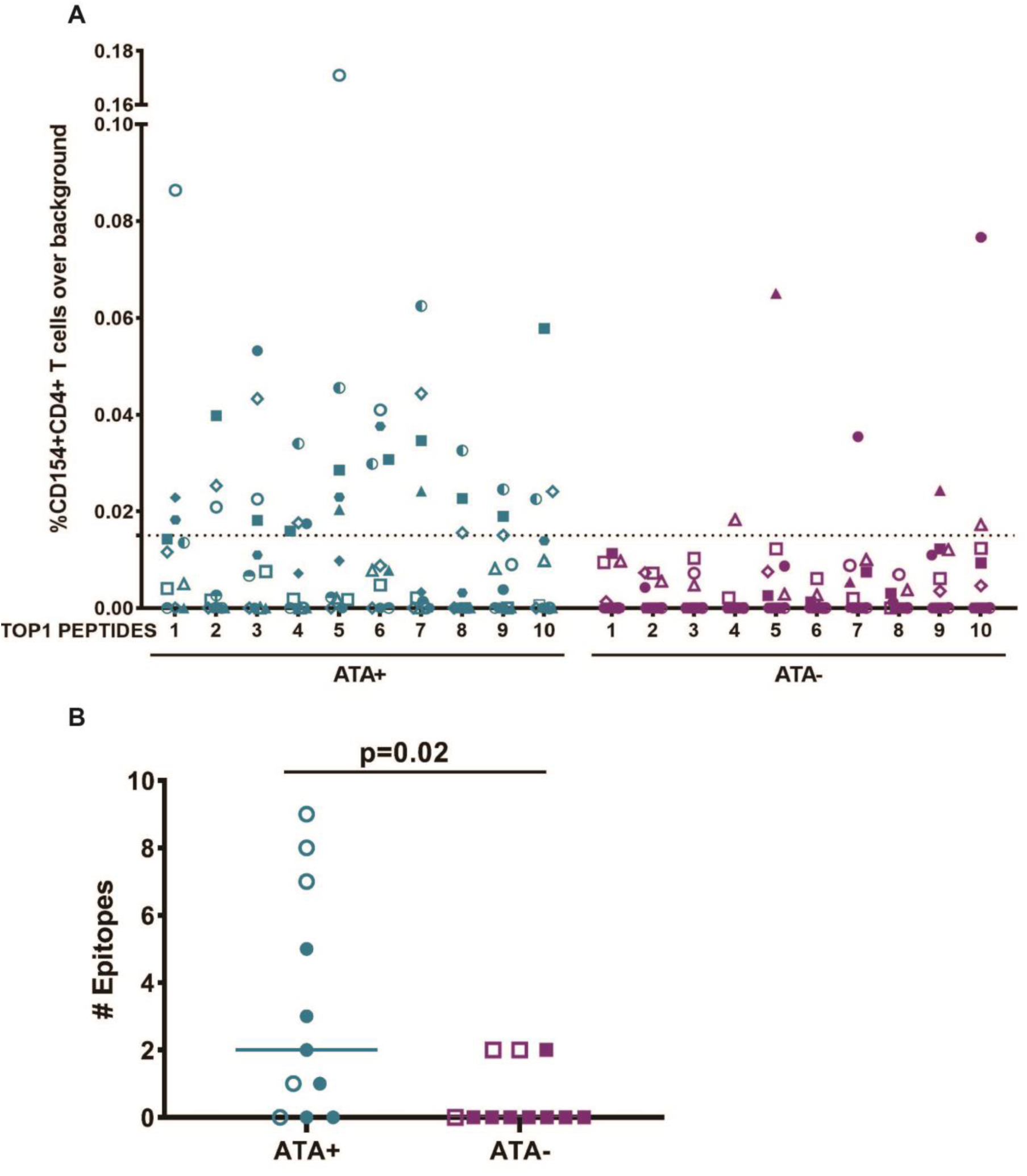
Distribution of CD4+ T cell responses to naturally presented TOP1 epitopes in scleroderma. **(A)** %CD154+CD4+ T cell responses over background observed with media alone for 11 ATA-positive and 11 ATA-negative scleroderma patients for each of the 10 naturally processed TOP1 peptides identified using NAPA are shown. The dotted line represents the cutoff for positivity that was defined as >95^th^ percentile of the T cell responses observed in ATA-negative patients (95^th^ percentile=0.015%). Each unique colored symbol represents a different patient. **(B)** The number of peptides eliciting a response in ATA-positive (n=11) and ATA-negative (n=11) scleroderma patients is shown (Mann-Whitney, p=0.02). A p-value <0.05 was considered significant throughout. Open symbols represent patients that possess at least one previously defined ATA-associated *HLA-DRB1* allele.

### Diversity of TOP1-specific T cell responses associated with ILD severity

Given the breadth of the TOP1-specific CD4+T cell response across multiple commonly presented epitopes and known association of TOP1-specific CD4+ T cell frequency with ILD(11), we examined whether the number of TOP1-epitopes recognized was associated with the severity of ILD in ATA-positive patients with scleroderma. We evaluated the lowest forced vital capacity (FVC) recorded during the course of clinical follow-up for each patient as a measure of the severity of lung fibrosis. We focused this analysis on the 9 ATA-positive patients with radiographically defined scleroderma-associated ILD. A significant negative association was observed between the minimum FVC and the number of TOP1 epitopes that elicited a T cell response (p=0.025; Fig. 7A). While this relationship was also observed in response to whole TOP1 protein (p=0.002; Fig. 7B), it is remarkable that these 10 commonly presented epitopes were sufficient to identify T cell responses associated with more severe lung disease. This implicates CD4+ T cells targeting commonly presented TOP1 epitopes in the pathogenesis of scleroderma-ILD.

**Fig. 7.**
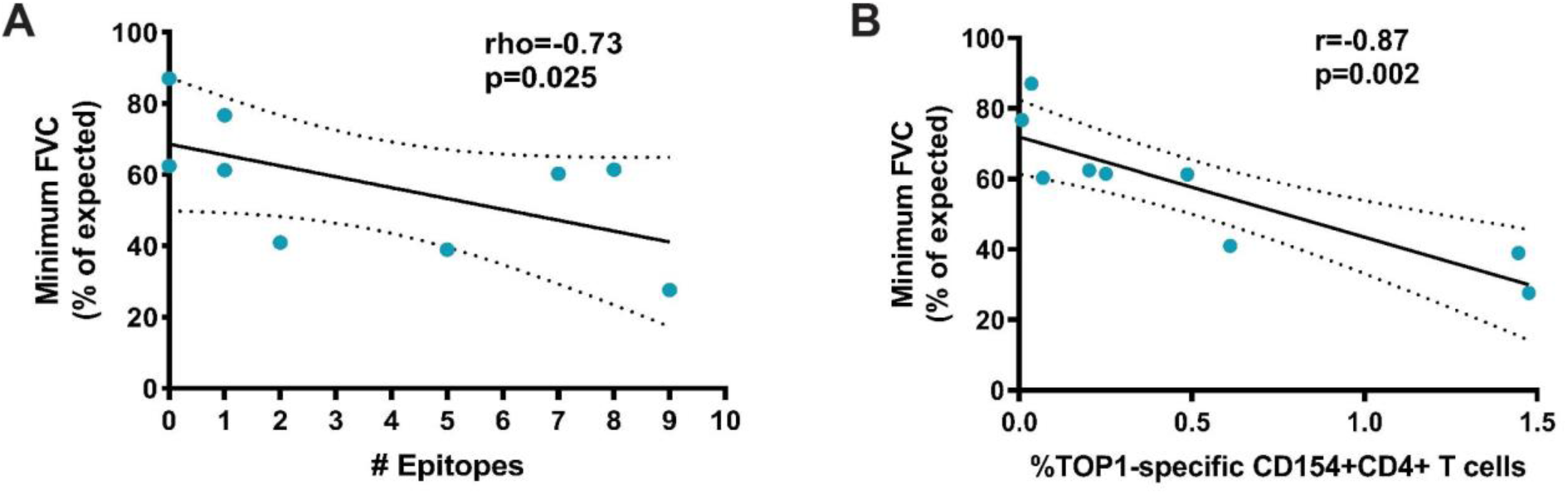
Clinical significance of CD4+ T cell responses to naturally presented TOP1 epitopes in scleroderma. The minimum FVC (% of expected) was plotted against **(A)** the number of TOP1 epitopes eliciting a T cell response and **(B)** the percentage of CD154+CD4+ T cells responses elicited against the whole TOP1 protein in ATA-positive scleroderma patients with clinically documented ILD (n=9). The correlation between minimum FVC and the number of stimulatory TOP1 epitopes was analyzed using a Spearman’s rho test, and the correlation with the magnitude of the T cell responses to the TOP1 protein was analyzed using a Pearson’s r test. A p-value <0.05 was considered significant throughout.

## Discussion

Our study supports a mechanism for how immunogenetically diverse patients affected by the autoimmune disease scleroderma can develop convergent immunologic and phenotypic characteristics. We have successfully developed and employed a robust method for the identification of biologically relevant naturally processed TOP1-specific CD4+ T cell autoantigenic epitopes leveraging antigen-presenting cells obtained directly from patients. Our data show that the presentation of a common set of TOP1 epitopes is shared by individuals with distinct HLA-DR variants and define how unique features present on both sides of the “peptide-HLA-DR interaction” synergize to drive this process. These findings, together with the observation that the breadth of the CD4+ T cell responses to these TOP1 epitopes is associated with presence and severity of lung fibrosis, offer novel insights into the pathogenesis of scleroderma.

Our study has shed light on one of the most proximal events in the stimulation of autoreactive TOP1-specific T cells, the presentation of autoantigenic epitopes by ATA-associated HLA-DR variants. We identified shared motifs in the peptide binding grooves of seemingly unrelated HLA-DR variants found in patients with ATA-positive scleroderma (^9^E-Y-S-T-S/G-E^14^ and ^67^F-L-E-D-R-A-A/L^74^) that enabled the presentation of a common set of TOP1 epitopes. Importantly, the ^67^F-L-E-D-R^71^ motif was previously identified by Kuwana *et al.* in 1993 and hypothesized to be a common feature of alleles associated with ATA in different ethnicities(18, 19, 24), but the mechanism for this was unknown. Our data provide direct evidence that the presence of HLA-DR alleles containing this or homologous sequences is associated with the amplified presentation of a higher number of TOP1 peptides as well as with autoreactive CD4+ T cell responses directed towards multiple TOP1 epitopes. The finding in scleroderma patients of shared motifs in seemingly diverse alleles is analogous to the “shared epitope hypothesis” proposed in subjects with rheumatoid arthritis (25, 26). The “shared epitope” is a conserved amino acid sequence (^70^R/Q-R/K-R-A-A^74^) present in the HLA-DRβ chain of a group of several HLA-DR variants that, collectively, exhibit a strong association with the development of rheumatoid arthritis (25). The shared epitope is hypothesized to lead to the presentation of common “arthritogenic” autoreactive peptides(27), though this has not yet been directly demonstrated. The analysis in RA patients of naturally processed antigenic peptides following our experimental approach (NAPA) may provide the evidence to confirm common arthritogenic peptide presentation in patients with shared epitope alleles. Although the collective analysis of alleles carrying a common motif has not yet been widely adopted for genetic studies in scleroderma, our data provides strong evidence for a shared epitope-like motif in scleroderma and bolsters the rationale for the use of such analyses in future studies.

The identification of the shared presentation of multiple TOP1 epitopes has also enabled the definition of two distinct groups of TOP1 peptides that differ in their interaction with HLA-DR. Group A TOP1 peptides contain a single core epitope that is bound in the same register to multiple HLA variants, likely interacting with elements in the peptide binding grove sharing similar features. Group B TOP1 peptides contain multiple distinct epitopes that utilize different anchor residues to bind to diverse HLA variants. While promiscuous peptides have been described in the setting of cancer, infection, and mouse models of autoimmunity (28–32), to our knowledge such peptides have not been described to be naturally presented in patients with autoimmunity. Further biophysical studies are required to define the precise anchor residues responsible for presentation of promiscuous TOP1 peptides in scleroderma, and to determine if this mechanism is operative in other autoimmune diseases.

Although the study of natural antigen processing by mo-DCs revealed a set of immunodominant TOP1 epitopes, this likely does not represent all of the TOP1 epitopes capable of driving pathogenic anti-TOP1 immune responses in scleroderma. There are several technical and biological explanations for this, including that NAPA may not have been able to capture TOP1 peptides that were presented at a low frequency or affinity, reflecting a finite sensitivity of the assay. Moreover, while dendritic cells are the most effective antigen presenting cells in activating T cell responses, other cells like macrophages and B lymphocytes express MHC class II processing machinery and have the ability to present CD4+ T cell epitopes(33). Therefore, additional TOP1 epitopes could be revealed by studying the processing of TOP1 by other antigen presenting cells. In addition, presentation of unique autoantigenic epitopes has been observed in resident antigen presenting cells present in disease target tissue in mouse models of diabetes, suggesting that presentation of tissue-specific CD4+ T cell epitopes may also occur(34, 35). Despite the above limitations, the discovery of immunorelevant TOP1 epitopes presented by mo-DCs and recognized by autoreactive T cells in the majority of patients with ATA-positive scleroderma, likely mirrors the critical role of dendritic cells in initiating a primary autoimmune response to TOP1.

The pathogenic relevance of the convergent presentation of TOP1 epitopes in patients with scleroderma is illustrated by the detection of TOP1-specific CD4+ T cell responses to this core set of epitopes in >70% of patients with ATA-positive scleroderma. We showed enrichment of TOP1-epitope-specific T cells in ATA-positive compared to ATA-negative patients, and a significant relationship between the number of TOP1 epitopes recognized and the severity of lung fibrosis. Unique HLA-DR variants have been significantly linked to scleroderma, and in particular, to specific disease manifestations such as ILD(36). Importantly, *HLA-DRB*1*11:01 has shown association with ILD only in the presence of ATA. In a British cohort(37), the relative risk of developing ILD in patients carrying *HLA-DRB*1*11:01 was 2.7 (statistically not significant), but this increased to 15.8 in subjects with both *HLA-DRB*1*11:01 and positive ATA. Together with the evidence provided by our study, this suggests a model where specific HLA variants are mechanistically linked to the development of scleroderma ILD by favoring the selection of TOP1-specific autoreactive T cells.

In our previous work, we have shown that the frequency of TOP1-specific CD4+ T cells is associated with the severity of lung fibrosis(11). This new study extends that observation by successfully defining the epitope-specificity of these autoreactive T cells. We demonstrate that a core set of 10 TOP1 epitopes elicits robust specific T cell responses in the majority of studied ATA positive scleroderma-ILD patients, with the breadth of epitopes recognized inversely associating with lung function. This has significant clinical implications and may inform the development of antigen-specific monitoring and therapeutic tools.

Tetramers made from MHC molecules bound to antigenic peptides can be used to identify, track, and study antigen-specific immune responses over time in peripheral blood, as well as target tissues (38). In scleroderma patients, TOP1 tetramers could therefore be a precise tool to study the frequency and phenotype of TOP1 specific T cells *ex vivo*. Longitudinal studies in early scleroderma cohorts will define whether the repertoire of TOP1 epitope-specific CD4+ T cells changes over time and predicts onset or progression of fibrotic lung disease. This is an important unmet need as ILD is among the leading causes of scleroderma-related mortality and no reliable biomarkers is clinically available to predict early on the development and the clinical trajectory of lung involvement(39). The core set of TOP1 epitopes identified in our investigation can also represent viable candidates for the development of novel antigen-specific therapies. By understanding the polyclonal nature of the autoimmune responses and the advances of medical bioengineering, we can ensure the delivery of biologically relevant epitopes using tools like nanoparticles or other carriers. Such therapies can induce the formation of antigen-specific regulatory networks and have shown promise in pre-clinical models of autoimmune disease and an increasing number of early phase clinical trials(40, 41).

In this work we successfully leveraged the natural antigen processing machinery of a patient’s own antigen presenting cells to define immunorelevant CD4+ T cell epitopes of the autoantigen TOP1. Our findings have implications for our understanding and conceptualization of the development of autoreactive immune responses in scleroderma and other autoimmune diseases. Rather than each individual presenting and recognizing a unique set of autoantigenic peptides, our data supports an alternative model in which a convergent set of self-epitopes is presented by diverse HLA-DR variants, leading to the development of similar autoimmune responses and clinical features.

## Materials and Methods

### Patient Characterization

Patients in this study were drawn from an observational cohort of patients with scleroderma followed at the Johns Hopkins Scleroderma Center. All patients participating in this study met the 2013 Classification Criteria for Systemic Sclerosis or met at least 3 out of 5 criteria for CREST (i.e. calcinosis, Raynaud’s phenomenon, esophageal dysmotility, sclerodactyly, telangiectasia)(42, 43). The Johns Hopkins institutional review board approved the study and all patients signed written informed consent.

Six ATA-positive patients were recruited for NAPA. For T cell stimulation assays, 11 patients with ATA and 11 patients with either anti-centromere antibodies (ACA) or anti-RNA polymerase 3 (ARA) antibodies, who were negative for ATA, were recruited for this study. Demographic data, scleroderma subtype (limited versus diffuse skin disease)(43), autoantibody specificities, disease duration from the time of first symptoms (Raynaud or non-Raynaud), smoking status (current), and medication history at the time of the blood draw, were obtained from the Scleroderma Center’s database. In order to capture the most developed disease phenotype, the lowest FVC from the pulmonary function test results were utilized. FVC was standardized to age and sex(44, 45). Any missing clinical information was obtained from a comprehensive review of the electronic medical records. Patients were classified as having limited cutaneous scleroderma, if scleroderma skin sclerosis was restricted to skin distal to the knees and elbows and/or the face, and diffuse cutaneous scleroderma, if skin sclerosis involved the chest, abdomen, and/or proximal extremities (i.e. thighs or upper arms)(43). The disease duration was calculated both from the first non-Raynaud’s disease related symptom as well as from the first occurrence of Raynaud’s phenomenon to the time of the serum sample processing. Gastrointestinal involvement was considered present if Medsger score was equal or greater than 1(46). ILD was defined based on evidence of pulmonary fibrosis on chest CT. Renal crisis was confirmed by renal biopsy in case of an acute increase in blood pressure. Current immunosuppression was recorded at the time of the blood sampling (prednisone, mycophenolate mofetil, methotrexate, cyclophosphamide, azathioprine, rituximab, tocilizumab). Clinical data are summarized in Table S4.

### Natural antigen processing assay (NAPA)

#### Monocyte-derived dendritic cell differentiation and antigen-loading

Peripheral blood mononuclear cells (PBMCs) were isolated by density-gradient centrifugation (Ficoll-Paque Plus, GE Healthcare) from 40ml of whole blood obtained from 6 ATA-positive patients. CD14 MicroBeads were used to positively select CD14+ monocytes and cells were differentiated for 7 days into monocyte-derived dendritic cells (mo-DCs), using Mo-DC Differentiation Medium (Miltenyi Biotec GmbH). Mo-DCs (2-5 × 10^6^) were reconstituted at 2x10^6^ cells/mL in Mo-DC Medium and incubated overnight at 37°C in 5% CO_2_ with 400 ug of custom-made human recombinant TOP1 protein purified from baculovirus-infected insect cells (GenScript Biotech Corp, Piscataway, NJ, USA). For control unpulsed experiments, mo-DCs were incubated overnight in media alone.

#### Isolation of HLA-DR-peptide complexes

Antigen-pulsed mo-DCs were collected and washed with cold buffer A (PBS, 2 mM EDTA; pH 7.4-7.6). The cells were then lysed in cold buffer B (1% CHAPS, 1 mM EDTA, protease inhibitor cocktail) and incubated at 4° C on a rocking table for 15 minutes. This lysis step was repeated, and insoluble material was removed by centrifugation at 14,000 x g for 15 minutes at 4° C. The supernatant containing the HLA-DR-peptide complexes was incubated with Protein G Dynabeads (Thermo Fisher Scientific) coated with 20 ug of anti-HLA-DR antibody (clone L243, BioLegend) at 4° C for one hour with inversion, per manufacturer’s instructions. For the isotype control experiment, we used IgG2α, κ isotype control antibody (LEAF Purified Mouse IgG2α, κ isotype control, Clone MOPC-173, BioLegend) in parallel samples for immunoprecipitation. The beads were then washed 4-times with lysis buffer B, and 3-times with buffer C (20 mM Tris, 150 mM NaCl, pH 7.4). After this extensive washing, the beads were incubated with 0.1% TFA for 1 minute at room temperature to elute the HLA-DR-bound peptides. The eluate was collected and incubation with 0.1% TFA was repeated. The pooled eluate was loaded onto a 10kD centrifugal filter (Microcon Centrifugal Filter Device, Merck Millipore) and centrifuged at 14,000 x g until all of the fluid passed through. The filter was washed with 0.1 % TFA and pooled flow-through was lyophilized by vacuum centrifugation.

#### Mass Spectrometry Analysis

Samples were reconstituted in 20 ul of 25% acetonitrile, 0.05% formic acid, and interfering reagents were removed by strong cation exchange (SCX) on a “stage tip”(47). Peptides were eluted from the SCX tip with 20 ul of 400mM ammonium bicarbonate in 25% acetonitrile and 0.05% formic acid and dried by vacuum centrifugation. Peptides were resuspended by adding 4ul of 10% acetonitrile, 0.1% formic acid, vortexing and diluting with 14 ul of 0.1% formic acid. The entire 18 ul sample was injected onto a 2 cm reverse phase trap (YMC ODS-A 10um particles, 120 Angstroms Kyoto, Japan) and separated on a 75 nm x 20 cm reverse phase column (Reprosil 3um particles, 100A pore size Ammerbuch-Entringen) using 2% to 90% acetonitrile gradient in 0.1% formic acid over 90 mins at 300 nl/min on an EasyLC nanoLC (Thermo Scientific) interfaced with Q-Exactive Plus mass spectrometer (Thermo Scientific). Eluting peptides were electrosprayed at 2kV, analyzed at 70K resolution (MS) and sequenced by collisional dissociation (HCD) with peptide fragments analyzed at 35K resolution (MS2). Ion target values for MS and MS2 were set at 3e6 and 1e5 with 100 and 150 milliseconds, respectively, and a normalized collision energy of 28. Data files were searched against the 2017 RefSeq83 human database (which contains the TOP1 protein sequence, NP_003277) using PEAKS 7 (Bioinformatic Solutions, Inc.) with the following criteria: 5 ppm mass tolerance for peptide masses; 0.02 daltons for peptide fragments; oxidation of methionine and deamidation of asparagine and glutamine as variable modifications. The database search was performed with no enzyme designation since no exogenous enzyme was added to produce the peptides. The search results were filtered at the 1% (false discovery rate) FDR level using the PEAKS decoy-fusion algorithm. Histograms were created to determine the length distribution of HLA-DR bound peptides, and peptides derived from keratin or dermcidin were excluded as likely artifactual contaminants.

### *HLA-DRB1* genotyping and sequence alignment

Whole blood was collected using PAXgene Blood DNA tubes and DNA was extracted according to manufacturer’s instructions (PreAnalytiX). High-resolution *HLA-DRB1* typing was performed by next generation sequencing at the Johns Hopkins University Immunogenetics Laboratory, as previously described(48). Amino acid sequences encoded for by the *HLA-DRB1* alleles of interest were obtained using the online IMGT/HLA submission tools provided by the Immune Polymorphism Database (IPD)(49, 50). Peptide contact and pocket residues in each HLA-DRβ1 chain were identified using the online MHC motif viewer(51). Sequence alignment and consensus sequence visualization was performed using Jalview version 2.10.5 software(52). Affinity and register of peptides for HLA-DR variants was predicted using the online NetMHCII 2.3 algorithm(22).

### Protein Structure Images

Linear mapping of patient-specific TOP1 epitopes was plotted using R (R Core Team (2018). R: A language and environment for statistical computing. R Foundation for Statistical Computing, Vienna, Austria. URL https://www.R-project.org/.). The three-dimensional location of TOP1 peptides identified using NAPA within the TOP1 protein crystal structure (1K4T) and location of consensus peptide binding motifs within HLA-DR11 (6CPN) was visualized using the PyMOL Molecular Graphics System, version 2.3 (Schrödinger, LLC).

### T cell stimulation

Fresh PBMCs from randomly selected ATA-positive (n=11) and ATA-negative patients (n=11) were isolated by Ficoll density-gradient centrifugation (Ficoll-Paque Plus, GE Healthcare). The cells were resuspended in RPMI medium supplemented with 5% autologous serum, transferred onto a 96-well plate (1.5 × 10^6^ cells/well) and cultured at 37 °C, as previously described(11, 23). Anti-human CD40 blocking antibody (1 ug/ml; G28.5, BioLegend) was added to the cells in order to prevent CD154 internalization. PBMCs were then stimulated for 18 hours using 2.5 uM of each candidate TOP1 peptide (Table 1), synthesized as free peptides with a purity >95%, or 1.45 ug/mL human recombinanct TOP1 protein, purified from baculovirus-infected insect cells (Genscript Biotech Corp). Following stimulation, cells were harvested, washed with PBS and stained with BV510-conjugated anti-CD3 (OKT3, BioLegend), Pacific Blue-conjugated anti-CD4 (RPA-T4, BD Pharmingen), APC-H7-conjugated anti-CD8 (SK1, BD Biosciencies), PE –conjugated anti-CD154 (TRAP1, BD Pharmingen), and Live/Dead Fixable Dead Cell Stain Kit (Molecular Probes). The percentage of live CD4+ T cells that upregulated CD154 were quantified by flow cytometry using a FACSAria (BD Biosciences) instrument at Bayview Immunomics Core Facility and analysis was conducted using FCS Express software, version 5 (De Novo Software). Pie charts were calculated by subtracting the sum of %CD154+CD4+ T cell responses to TOP1 peptides defined as Group A peptides (TOP1 peptides 1, 2, 8, 9 and 10), and the sum of the responses to Group B peptides (TOP1 peptides 3, 4, 5, 6 and 7) from the %CD154+CD4+ T cell responses to whole TOP1 protein. If the subtraction was >0, the remaining sum was defined as other peptides.

### Statistical analyses

The naturally presented peptides were analyzed using the PANTHER 14.1 gene list analysis tool statistical overrepresentation test for “biological processes”. This was performed to determine if the naturally processed peptides were derived from proteins involved in specific processes and too identify those biological processes that were over-represented relative to the human genome(53, 54). The “GO biological processes complete” annotation set was queried and the FDR and Fisher’s exact p-value for each pathway was calculated. The median number of peptides eliciting positive responses in ATA-positive versus ATA-negative patients was compared using a Mann-Whitney test. A cutoff of greater than the 95^th^ percentile of the responses observed amongst the ATA-negative patients (%CD154+CD4+Tcells > 0.015%) was used to define a positive CD4+ T cell response to antigen stimulation. The association between minimum FVC and the number of stimulatory TOP1 epitopes was analyzed using the Spearman’s rho test. The association between minimum FCV and the percentage of TOP1-specific CD154+CD4+ T cells was analyzed using the Pearson’s r. Statistical analyses were performed using Prism version 7.03 (Graphpad) and Stata version 14.2 (Stata Corp) software. Statistical significance was defined as a two-sided p-value ≤0.05 throughout.

## Acknowledgments

We would like to thank the research coordinators (Adrianne Woods, Margaret Sampedro, and Gwendolyn Leatherman), physicians and rheumatology fellows of the Johns Hopkins Scleroderma Center for their support in obtaining the patient samples and data; the patients of the Johns Hopkins Scleroderma Center who participated in this study; Scheherazade Sadegh-Nasseri, Ph.D. for critical review of the data; and Felipe Andrade M.D., Ph.D. for critical review of the manuscript.

## Author contributions

E.T, A.F, F. B. and E.D designed the study and guided data interpretation. E.T. and E.D lead data acquisition and analysis. T.G. contributed to assay development and data acquisition. R.N.C guided mass spectrometry and troubleshooting; R.N.O. performed mass spectrometry and analyzed the data. Z.H.M. and A.A.S. performed the statistical analysis of clinical variables. Z.H.M, A.A.S., and F.M.W. provided clinical samples and data. All authors contributed to the review of the manuscript and approved the final version for publication.

## Data and materials availability

All data associated with this study are present in the paper or the Supplementary Materials.

## Supplementary Materials

**Table S1.** Mass Spectrometry Analysis of peptides eluted from mo-DCs of 6 ATA-positive Scleroderma patients using anti-HLA-DR antibody to immunoprecipitate the HLA-DR:peptide complexes (Excel file).

**Table S2.** Mass Spectrometry Analysis of peptides eluted from unpulsed mo-DCs using anti-HLA-DR antibody to immunoprecipitate the HLA-DR:peptide complexes (Excel file).

**Table S3.** Mass Spectrometry Analysis of peptides eluted from mo-DCs using anti-HLA-DR or isotype control antibody to immunoprecipitate the HLA-DR:peptide complexes (Excel file).

**Table S4.** Demographic and clinical characteristics of the ATA-positive and ATA-negative SSc patients included in T cell stimulation assays.

| Variable | ATA-positive (n=11) | ATA-negative (n=11) |
| --- | --- | --- |
| Age at sampling, means ± SD (years) | 56.8 ± 13.2 | 58.8 ± 14.8 |
| Female, no (%) | 10 (90.9) | 11 (100) |
| Race, no (%) |  |  |
| Caucasian | 7 (63.6) | 11 (100) |
| Black | 3 (27.3) | 0 (0) |
| Asian | 1 (9.1) | 0 (0) |
| Smoking status (current), no (%) | 3 (27.3) | 2 (18.2) |
| Scleroderma type, diffuse, no (%) | 6 (54.5) | 2 (18.2) |
| SSc duration |  |  |
| RP onset, means ± SD (years) | 18.7 ± 7.4 | 15.8 ± 11.8 |
| 1st non-RP symptom, means ± SD (years) | 6.5 ± 3.6 | 10.7 ± 11.0 |
| Interstitial Lung Disease, no (%) | 9 (81.8) | 0 (0) |
| Gastrointestinal Involvement, no (%) | 11 (100) | 10 (90.9) |
| Renal Crisis, no (%) | 0 (0) | 0 (0) |
| Autoantibodies |  |  |
| Anti-topoisomerase | 11 (100) | 0 (0) |
| Anti-centromere | 0 (0) | 10 (90.9) |
| Anti-RNA polymerase | 0 (0) | 1 (9.1) |
| Current Immunosuppression, no (%) | 7 (63.6) | 2 (18.2) |

\*ATA= anti-Topoisomerase-I antibody, SSc= systemic sclerosis, RP=Raynaud phenomenon, RVSP= right ventricular systolic pressure, SD=standard deviation, no=number

**Table S5.** CD4+ T cell responses of ATA-positive and ATA-negative scleroderma patients to TOP1 peptides identified using NAPA.

| TOP1 Peptide |  |  |  |  |  |  |  |  |  |  |  |  |  |
| --- | --- | --- | --- | --- | --- | --- | --- | --- | --- | --- | --- | --- | --- |
| ID | HLA-DRβ1 |  | 1 | 2 | 3 | 4 | 5 | 6 | 7 | 8 | 9 | 10 | #Responses/<br>Patient |
| ATA-positive |  |  |  |  |  |  |  |  |  |  |  |  |  |
| FW-1990 | 08:04 | 13:01 | - | + | + | + | + | + | + | + | + | + | 9 |
| FW-1963 | 01:02 | 11:01 | - | - | + | + | + | + | + | + | + | + | 8 |
| FW-0592 | 15:01 | 11:01 | - | + | + | + | - | - | + | + | + | + | 7 |
| FW-1858 | 11:03 | 13:01 | + | + | + | - | + | + | - | - | - | - | 5 |
| FW-1994 | 03:01 | 04:01 | + | - | - | - | + | + | - | - | - | - | 3 |
| FW-2235 | 01:01 | 01:01 | - | - | - | - | + | - | + | - | - | - | 2 |
| FW-2147 | 15:03 | 08:04 | - | - | - | + | - | - | - | - | - | - | 1 |
| FW-2337 | 15:03 | 09:01 | + | - | - | - | - | - | - | - | - | - | 1 |
| FW-1874 | 01:01 | 15:01 | - | - | - | - | - | - | - | - | - | - | 0 |
| FW-2019 | 11:04 | 11:04 | - | - | - | - | - | - | - | - | - | - | 0 |
| FW-2375 | 01:01 | 13:01 | - | - | - | - | - | - | - | - | - | - | 0 |
| #Responses/Peptide |  |  | 3 | 3 | 4 | 4 | 5 | 4 | 4 | 3 | 3 | 3 |  |
| ATA-negative |  |  |  |  |  |  |  |  |  |  |  |  |  |
| FW-2464 | 11:01 | 04:07 | - | - | - | - | - | - | + | - | - | + | 2 |
| FW-0896 | 08:01 | 11:04 | - | - | - | - | + | - | - | - | + | - | 2 |
| FW-0620 | 08:01 | 11:03 | - | - | - | + | - | - | - | - | - | + | 2 |
| FW-2170 | 03:01 | 07:01 | - | - | - | - | - | - | - | - | - | - | 0 |
| FW-2124 | 03:01 | 04:07 | - | - | - | - | - | - | - | - | - | - | 0 |
| FW-2301 | 01:01 | 03:01 | - | - | - | - | - | - | - | - | - | - | 0 |
| FW-2470 | 03:01 | 04:04 | - | - | - | - | - | - | - | - | - | - | 0 |
| FW-2119 | 08:01 | 04:07 | - | - | - | - | - | - | - | - | - | - | 0 |
| FW-1863 | 04:07 | 04:07 | - | - | - | - | - | - | - | - | - | - | 0 |
| FW-2398 | 11:04 | 11:04 | - | - | - | - | - | - | - | - | - | - | 0 |
| FW-2240 | 01:02 | 08:01 | - | - | - | - | - | - | - | - | - | - | 0 |
| # Responses/Peptide |  |  | 0 | 0 | 0 | 1 | 1 | 0 | 1 | 0 | 1 | 2 |  |

+= %CD154+CD4+ T cells above the 95th-percentile of ATA-negative patients; -= %CD154+CD4+ T cells below the 95th-percentile of ATA-negative patients

**Fig. S1.**
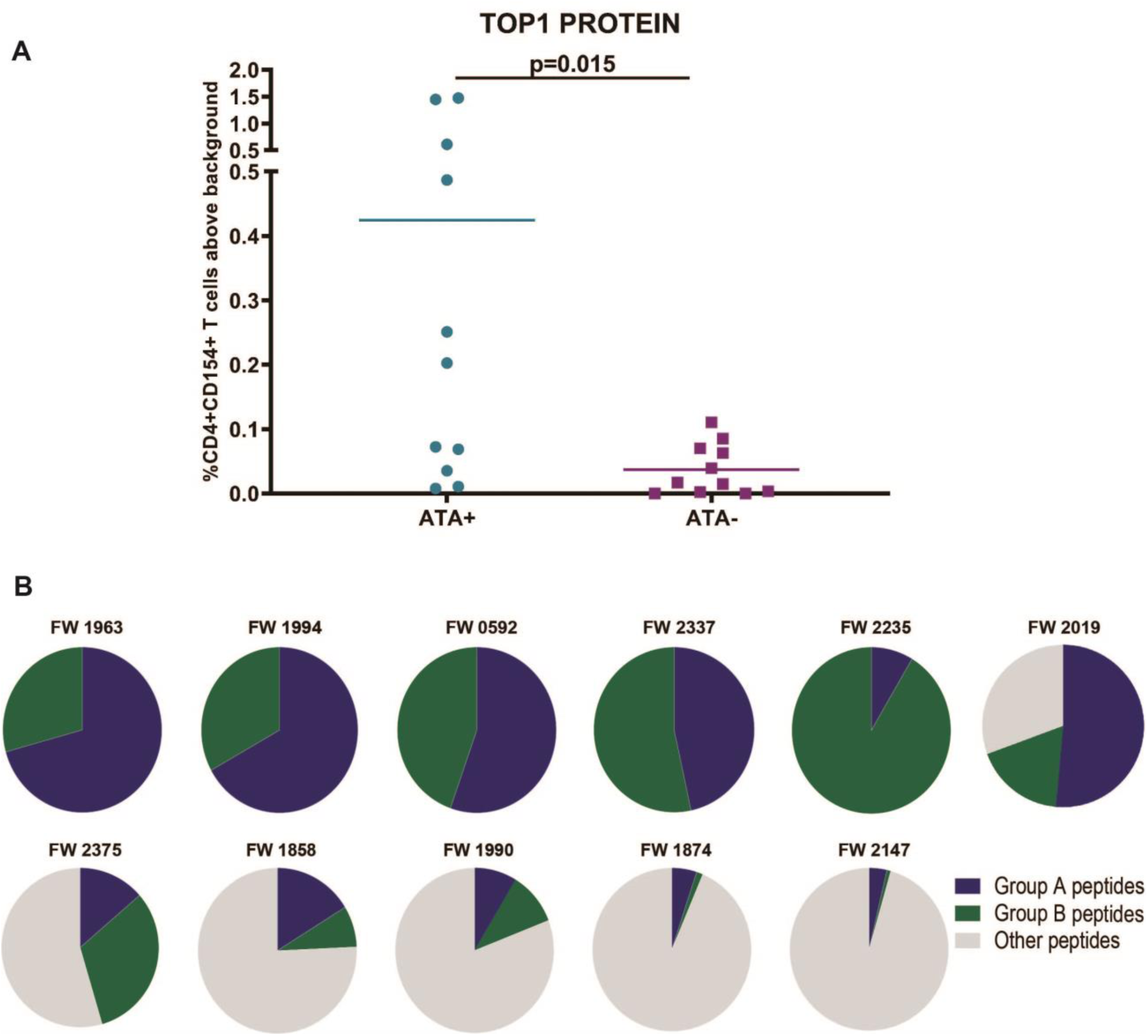
TOP1 peptide-specific T cell responses compared to T cell responses to whole TOP1 protein. **(A)** Frequency of CD154+CD4+ T cells against whole TOP1 molecule between 11 ATA-positive and 11 ATA-negative SSc patients. **(B)**The proportion CD154+CD4+ T cell responses against Group A TOP1 peptides (dark blue) or Group B TOP1 peptides (dark green) out of the total CD154+CD4 T cell response to whole TOP1 protein (grey) is shown for each patient.

## References and Notes

1. S.-H. Yang, et al., The molecular basis of immune regulation in autoimmunity. Clinical Science 132, 43–67 (2018).

2. P. Brodin, M. M. Davis, Human immune system variation. Nat. Rev. Immunol. 17, 21–29 (2017).

3. E. W. Newell, M. M. Davis, Beyond model antigens: high-dimensional methods for the analysis of antigen-specific T cells. Nat. Biotechnol. 32, 149–157 (2014).

4. A. Gabrielli, E. V. Avvedimento, T. Krieg, Scleroderma. New England Journal of Medicine 360, 1989–2003 (2009).

5. F. Boin, A. Rosen, Autoimmunity in systemic sclerosis: current concepts. Curr Rheumatol Rep 9, 165–172 (2007).

6. U. A. Walker, et al., Clinical risk assessment of organ manifestations in systemic sclerosis: a report from the EULAR Scleroderma Trials And Research group database. Ann. Rheum. Dis. 66, 754–763 (2007).

7. M. Kuwana, T. A. Medsger, T. M. Wright, T cell proliferative response induced by DNA topoisomerase I in patients with systemic sclerosis and healthy donors. J. Clin. Invest. 96, 586–596 (1995).

8. M. Kuwana, C. A. Feghali, T. A. Medsger, T. M. Wright, Autoreactive T cells to topoisomerase I in monozygotic twins discordant for systemic sclerosis. Arthritis Rheum. 44, 1654–1659 (2001).

9. S. Veeraraghavan, et al., Mapping of the immunodominant T cell epitopes of the protein topoisomerase I. Ann. Rheum. Dis. 63, 982–987 (2004).

10. P. Q. Hu, J. J. Oppenheim, T. A. Medsger, T. M. Wright, T cell lines from systemic sclerosis patients and healthy controls recognize multiple epitopes on DNA topoisomerase I. J. Autoimmun. 26, 258–267 (2006).

11. A. Fava, et al., Frequency of circulating topoisomerase-I-specific CD4 T cells predicts presence and progression of interstitial lung disease in scleroderma. Arthritis Res. Ther. 18, 99 (2016).

12. E. Appella, E. A. Padlan, D. F. Hunt, Analysis of the structure of naturally processed peptides bound by class I and class II major histocompatibility complex molecules. EXS 73, 105–119 (1995).

13. A. J. Godkin, et al., Naturally processed HLA class II peptides reveal highly conserved immunogenic flanking region sequence preferences that reflect antigen processing rather than peptide-MHC interactions. J. Immunol. 166, 6720–6727 (2001).

14. E. Darrah, et al., Proteolysis by Granzyme B Enhances Presentation of Autoantigenic Peptidylarginine Deiminase 4 Epitopes in Rheumatoid Arthritis. J. Proteome Res. 16, 355–365 (2017).

15. D. Mirano-Bascos, M. Tary-Lehmann, S. J. Landry, Antigen structure influences helper T-cell epitope dominance in the human immune response to HIV envelope glycoprotein gp120. Eur. J. Immunol. 38, 1231–1237 (2008).

16. P. Chairta, P. Nicolaou, K. Christodoulou, Genomic and genetic studies of systemic sclerosis: A systematic review. Hum. Immunol. 78, 153–165 (2017).

17. F. C. Arnett, et al., Major histocompatibility complex (MHC) class II alleles, haplotypes and epitopes which confer susceptibility or protection in systemic sclerosis: analyses in 1300 Caucasian, African-American and Hispanic cases and 1000 controls. Ann. Rheum. Dis. 69, 822–827 (2010).

18. F. Takeuchi, et al., Association of HLA-DR with progressive systemic sclerosis in Japanese. J. Rheumatol. 21, 857–863 (1994).

19. S. H. Kang, et al., Association of HLA class II genes with systemic sclerosis in Koreans. J. Rheumatol. 28, 1577–1583 (2001).

20. M. Kuwana, J. Kaburaki, Y. Okano, H. Inoko, K. Tsuji, The HLA-DR and DQ genes control the autoimmune response to DNA topoisomerase I in systemic sclerosis (scleroderma). J. Clin. Invest. 92, 1296–1301 (1993).

21. D. D. Gladman, et al., HLA markers for susceptibility and expression in scleroderma. J. Rheumatol. 32, 1481–1487 (2005).

22. K. K. Jensen, et al., Improved methods for predicting peptide binding affinity to MHC class II molecules. Immunology 154, 394–406 (2018).

23. C. G. Joseph, et al., Association of the autoimmune disease scleroderma with an immunologic response to cancer. Science 343, 152–157 (2014).

24. H. Furukawa, et al., Human Leukocyte Antigen and Systemic Sclerosis in Japanese: The Sign of the Four Independent Protective Alleles, DRB1*13:02, DRB1*14:06, DQB1*03:01, and DPB1*02:01. PLoS ONE 11, e0154255 (2016).

25. P. K. Gregersen, J. Silver, R. J. Winchester, The shared epitope hypothesis. An approach to understanding the molecular genetics of susceptibility to rheumatoid arthritis. Arthritis Rheum. 30, 1205–1213 (1987).

26. D. E. de Almeida, S. Ling, J. Holoshitz, New insights into the functional role of the rheumatoid arthritis shared epitope. FEBS Lett. 585, 3619–3626 (2011).

27. K. W. Wucherpfennig, J. L. Strominger, Selective binding of self peptides to disease-associated major histocompatibility complex (MHC) molecules: a mechanism for MHC-linked susceptibility to human autoimmune diseases. J. Exp. Med. 181, 1597–1601 (1995).

28. P. Panina-Bordignon, et al., Universally immunogenic T cell epitopes: promiscuous binding to human MHC class II and promiscuous recognition by T cells. Eur. J. Immunol. 19, 2237–2242 (1989).

29. J. Yang, et al., Autoreactive T cells specific for insulin B:11-23 recognize a low-affinity peptide register in human subjects with autoimmune diabetes. Proc. Natl. Acad. Sci. U.S.A. 111, 14840–14845 (2014).

30. A. L. Corper, et al., A structural framework for deciphering the link between I-Ag7 and autoimmune diabetes. Science 288, 505–511 (2000).

31. Y. Shi, et al., Promiscuous presentation and recognition of nucleosomal autoepitopes in lupus: role of autoimmune T cell receptor alpha chain. J. Exp. Med. 187, 367–378 (1998).

32. A. Seamons, et al., Competition between two MHC binding registers in a single peptide processed from myelin basic protein influences tolerance and susceptibility to autoimmunity. J. Exp. Med. 197, 1391–1397 (2003).

33. P. A. Roche, K. Furuta, The ins and outs of MHC class II-mediated antigen processing and presentation. Nat. Rev. Immunol. 15, 203–216 (2015).

34. N. J. Viner, C. A. Nelson, B. Deck, E. R. Unanue, Complexes generated by the binding of free peptides to class II MHC molecules are antigenically diverse compared with those generated by intracellular processing. J. Immunol. 156, 2365–2368 (1996).

35. J. F. Mohan, et al., Unique autoreactive T cells recognize insulin peptides generated within the islets of Langerhans in autoimmune diabetes. Nat. Immunol. 11, 350–354 (2010).

36. C. J. W. Stock, E. A. Renzoni, Genetic predictors of systemic sclerosis-associated interstitial lung disease: a review of recent literature. Eur. J. Hum. Genet. 26, 765–777 (2018).

37. G. C. Fanning, K. I. Welsh, C. Bunn, R. Du Bois, C. M. Black, HLA associations in three mutually exclusive autoantibody subgroups in UK systemic sclerosis patients. Br. J. Rheumatol. 37, 201–207 (1998).

38. J. D. Altman, M. M. Davis, MHC-Peptide Tetramers to Visualize Antigen-Specific T Cells. Curr Protoc Immunol 115, 17.3.1–17.3.44 (2016).

39. V. Cottin, K. K. Brown, Interstitial lung disease associated with systemic sclerosis (SSc-ILD). Respir. Res. 20, 13 (2019).

40. X. Clemente-Casares, et al., Expanding antigen-specific regulatory networks to treat autoimmunity. Nature 530, 434–440 (2016).

41. P. Serra, P. Santamaria, Antigen-specific therapeutic approaches for autoimmunity. Nat. Biotechnol. 37, 238–251 (2019).

42. F. van den Hoogen, et al., 2013 classification criteria for systemic sclerosis: an American college of rheumatology/European league against rheumatism collaborative initiative. Ann. Rheum. Dis. 72, 1747–1755 (2013).

43. E. C. LeRoy, T. A. Medsger, Criteria for the classification of early systemic sclerosis. J. Rheumatol. 28, 1573–1576 (2001).

44. R. J. Knudson, W. T. Kaltenborn, D. E. Knudson, B. Burrows, The single-breath carbon monoxide diffusing capacity. Reference equations derived from a healthy nonsmoking population and effects of hematocrit. Am. Rev. Respir. Dis. 135, 805–811 (1987).

45. J. L. Hankinson, J. R. Odencrantz, K. B. Fedan, Spirometric reference values from a sample of the general U.S. population. Am. J. Respir. Crit. Care Med. 159, 179–187 (1999).

46. T. A. Medsger, et al., A disease severity scale for systemic sclerosis: development and testing. J. Rheumatol. 26, 2159–2167 (1999).

47. J. Rappsilber, M. Mann, Y. Ishihama, Protocol for micro-purification, enrichment, pre-fractionation and storage of peptides for proteomics using StageTips. Nat Protoc 2, 1896–1906 (2007).

48. L. C. Cappelli, M. F. Konig, A. C. Gelber, C. O. Bingham, E. Darrah, Smoking is not linked to the development of anti-peptidylarginine deiminase 4 autoantibodies in rheumatoid arthritis. Arthritis Res. Ther. 20, 59 (2018).

49. J. Robinson, et al., The IPD and IMGT/HLA database: allele variant databases. Nucleic Acids Res. 43, D423–431 (2015).

50. J. Robinson, A. Malik, P. Parham, J. G. Bodmer, S. G. Marsh, IMGT/HLA database--a sequence database for the human major histocompatibility complex. Tissue Antigens 55, 280–287 (2000).

51. N. Rapin, I. Hoof, O. Lund, M. Nielsen, MHC motif viewer. Immunogenetics 60, 759–765 (2008).

52. A. M. Waterhouse, J. B. Procter, D. M. A. Martin, M. Clamp, G. J. Barton, Jalview Version 2--a multiple sequence alignment editor and analysis workbench. Bioinformatics 25, 1189–1191 (2009).

53. H. Mi, P. Thomas, “PANTHER Pathway: An Ontology-Based Pathway Database Coupled with Data Analysis Tools” in Protein Networks and Pathway Analysis, Y. Nikolsky, J. Bryant, Eds. (Humana Press, 2009), pp. 123–140.

54. H. Mi, et al., Protocol Update for large-scale genome and gene function analysis with the PANTHER classification system (v.14.0). Nature Protocols 14, 703–721 (2019).

